# Phenome-wide Heritability Analysis of the UK Biobank

**DOI:** 10.1101/070177

**Authors:** Tian Ge, Chia-Yen Chen, Benjamin M. Neale, Mert R. Sabuncu, Jordan W. Smoller

## Abstract

Heritability estimation provides important information about the relative contribution of genetic and environmental factors to phenotypic variation, and provides an upper bound for the utility of genetic risk prediction models. Recent technological and statistical advances have enabled the estimation of additive heritability attributable to common genetic variants (SNP heritability) across a broad phenotypic spectrum. However, assessing the comparative heritability of multiple traits estimated in different cohorts may be misleading due to the population-specific nature of heritability. Here we report the SNP heritability for 551 complex traits derived from the large-scale, population-based UK Biobank, comprising both quantitative phenotypes and disease codes, and examine the moderating effect of three major demographic variables (age, sex and socioeconomic status) on the heritability estimates. Our study represents the first comprehensive phenome-wide heritability analysis in the UK Biobank, and underscores the importance of considering population characteristics in comparing and interpreting heritability.

## Introduction

The heritability of a trait refers to the proportion of phenotypic variance that is attributable to genetic variation among individuals. Heritability is commonly measured as either the contribution of total genetic variation (broad-sense heritability, *H*^2^), or the fraction due to additive genetic variation (narrow-sense heritability, *h*^2^) (Visscher et al., 2008). A large body of evidence from twin studies has documented that essentially all human complex traits are heritable. For example, a recent meta-analysis of virtually all twin studies published between 1958 and 2012, encompassing 17,804 traits, reported that the overall narrow-sense heritability estimate across all human traits was 49%, although estimates varied widely across phenotypic domains (Polderman et al., 2015). Over the past decade, the availability of genome-wide genotyping has enabled the direct estimation of additive heritability attributable to common genetic variation (“SNP heritability” or 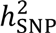) (Lee et al., 2011; Yang et al., 2010, 2011). These estimates provide a lower bound for narrow-sense heritability because they do not capture non-additive genetic effects such as dominance or epistasis, and contributions (e.g., from rare variants) that are not assayed by most genotyping microarrays and are not well tagged by genotyped variants. Nevertheless, estimates of SNP heritability can provide important information about the genetic basis of complex traits such as the proportion of phenotypic variation that could be explained by common-variant genome-wide association studies (GWAS).

However, heritability is not a fixed property of a phenotype but depends on the population in which it is estimated. As a ratio of variances, it can vary with population-specific differences in both genetic background and environmental variation (Visscher et al., 2008). For example, twin data have documented variations in the heritability of childhood IQ by socioeconomic status (Turkheimer et al., 2003), highlighting that different environment may have different relative contributions to the variance of a phenotype. In addition, heritability estimates for a range of complex phenotypes have been shown to vary according to the sex and age distributions of the sampled populations (Polderman et al., 2015). Identifying variables that may affect the heritability of complex traits has implications for the design of GWAS, highlighting subgroups and environmental conditions in which common-variant contributions may be diminished or magnified. To date, however, studies of complex trait heritability and the effect of modifying variables have largely examined individual phenotypes. Assessing the comparative heritability of traits estimated in independent samples may be misleading because of population-specific differences in genetic and environmental variance that may be operating in different cohorts.

The UK Biobank (http://www.ukbiobank.ac.uk) provides a unique opportunity to estimate and compare the heritability of traits across a broad phenotypic spectrum in a single population sample. The UK Biobank is a large prospective population-based cohort study that enrolled 500,000 participants aged 40-69 years between 2006 and 2010 (Sudlow et al., 2015). The study has collected a wealth of phenotypic data from questionnaires, physical and biological measurements, and electronic health records as well as genome-wide genotype data. Here we used a computationally efficient approach to estimating heritability for 551 complex traits, comprising both quantitative phenotypes and disease categories. We then examined how heritability estimates are modified by three major demographic variables: age, sex and socioeconomic status (SES). Our results underscore the importance of considering population characteristics in comparing heritability, and may inform efforts to apply genetic risk prediction models for a broad range of human phenotypes.

## Material and Methods

### Participants and Data Sources

This study utilized data from the baseline assessment of the UK Biobank, a prospective cohort study of 500,000 individuals (age 40-69 years) recruited across Great Britain during 2006-2010 (Sudlow et al., 2015). The protocol was approved by the Research Ethics Committee. The UK Biobank collected phenotypic data from a variety of sources including questionnaires regarding mental and physical health, food intake, family history and lifestyle, a baseline physical assessment, computerized cognitive testing, linkage with health records, and blood samples for biochemical and DNA analysis. Details about the UK Biobank project are provided at http://www.ukbiobank.ac.uk. Data for the current analyses were obtained under an approved data request (Ref: 13905).

### Genotyping and Quality Control

The interim release of the genotype data for the UK Biobank comprises 152,736 samples. Two closely related arrays from Affymetrix, the UK BiLEVE Axiom array and the UK Biobank Axiom array, were used to genotype approximately 800,000 markers with good genome-wide coverage. Details of the design of the arrays and sample processing can be found at http://biobank.ctsu.ox.ac.uk/crystal/refer.cgi?id=146640 and http://biobank.ctsu.ox.ac.uk/crystal/refer.cgi?id=155583.

Prior to the release of the genotype data, stringent quality control (QC) was performed at the Wellcome Trust Centre for Human Genetics, Oxford, UK. Procedures were documented in detail at http://biobank.ctsu.ox.ac.uk/crystal/refer.cgi?id=155580. We leveraged the QC metrics and removed samples that had mismatch between genetically inferred sex and self-reported sex, samples that had high genotype missingness or extreme heterozygosity not explained by mixed ancestry or increased levels of marriage between close relatives, and one individual from each pair of the samples that were 3^rd^ degree or more closely related relatives. We restricted our analysis to subjects that were self-reported white British and confirmed by principal component analysis (PCA) to be Caucasians. We further filtered out genetic markers that had high missing rate (>1%), low minor allele frequency (<1%), significant deviation from Hardy-Weinberg equilibrium (*p*<1e-7), and subjects that had high missing genotype rate (>1%). 108,158 subjects (age 40-73 years; female 52.84%) and 486,175 SNPs remained for analysis after QC. Supplementary Figure 1 shows the age distribution of the subjects that passed QC. The genetic similarity matrix was computed using all genotyped autosomal SNPs. All genetic analyses were performed using PLINK 1.9 (https://www.cog-genomics.org/plink2) (Chang et al., 2015).

### Phenotypic Variables

We analyzed every trait available to us that had a sufficient sample size to produce a heritability estimate with its standard error smaller than 0.1. The traits can be classified into the following 11 domains: cognitive functions, early life factors, health and medical history, life style, physical measures, psychosocial factors, sex-specific factors and sociodemographics. For continuous traits, we excluded samples that were more than 5 standard deviations away from the population mean to avoid extreme outliers and data recording errors. We only analyzed binary traits that had prevalence greater than 1% in the sample, so that we had enough power to get reliable heritability estimates. We typically binarized categorical variables at a meaningful threshold close to the median and then analyzed them as binary traits. For the specific cutoff-points used to binarize each categorical variable, see Supplementary Table 1.

We also analyzed a large number of self-reported illness codes and hospital in-patient diagnosis codes. Self-reported cancer and non-cancer illness codes were obtained through a verbal interview by a trained nurse at the UK Biobank assessment center on past and current medical conditions. Hospital in-patient diagnoses were obtained through medical records and were coded according to the International Classification of Diseases version-10 (ICD-10). Disease codes for each domain (self-reported cancer, self-reported noncancer illness, and ICD-10) were organized in a hierarchical tree structure; codes closer to the root of the tree are often less specific and have larger prevalence, while codes closer to the leaves are more specific but have lower prevalence. We analyzed every disease code that had prevalence greater than 1% in the sample. We also employed a data-driven approach to determine if a disease is sex-specific. More specifically, if the sample prevalence of a disease in males was more than 100 times larger than the sample prevalence in females, we defined the disease as male-specific and the analysis was restricted to males. The same approach was used to find female-specific diseases. See Supplementary Table 2 for all the disease codes we analyzed.

### Heritability Estimation

We consider the linear random effect model ***y*** = ***g*** + ***e***, where an *N*-dimensional trait *y* is partitioned into the sum of additive genetic effects *g* and unique (subject-specific) environmental effects *e*. The covariance structure of *y* is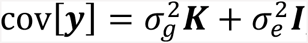, where *K* is the empirical genetic similarity matrix for each pair of individuals estimated from genome-wide SNP data (Yang et al., 2010, 2011), ***I*** is an identity matrix,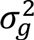 and 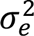 are the total additive genetic variance captured by genotyped common SNPs and the variance of unique environmental factors across individuals, respectively. SNP heritability is then defined as 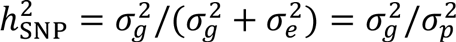, which measures the total phenotypic variance 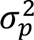 that can be explained by total additive genetic variance tagged by genotyped SNPs, and is a lower bound for the narrow-sense heritability *h*^2^. When covariates need to be incorporated into the model, i.e., ***y*** = ***Xβ*** + ***g*** + ***e***, where ***X*** is an *N*×*q* covariate matrix and ***β*** is a vector of fixed effects, an *N×(N – ą*) matrix *U* always exists, which satisfies ***U^T^U*** = ***I***, ***UU^T^*** = ***P*_0_**, ***U^T^X*** = 0, and ***P*_0_** = ***I*** – ***X*(*X^T^X***)^-1^ ***X^T^***. Applying *U^T^* to both sides of the model removes the covariate matrix (Ge et al., 2015).

To obtain unbiased estimates of 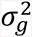 and 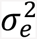, we used a computationally efficient, moment-matching approach, which is equivalent to the Haseman-Elston regression (Elston et al., 2000; Haseman and Elston, 1972; Sham and Purcell, 2001) and phenotype-correlation genetic-correlation (PCGC) regression (Golan et al.2014), and closely related to the LD score regression (Bulik-Sullivan et al., 2015b; Bulik-Sullivan 2015). Specifically, we regress the empirical estimate of the phenotypic covariance onto the matrices ***K*** and ***I***: 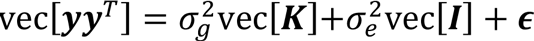, where vec[·] is the matrix vectorization operator that converts a matrix into a vector by stacking its columns, and **∊** is the residual of the regression. The ordinary least squares (OLS) estimator of this multiple regression problem can be explicitly written as 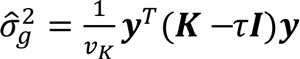 and 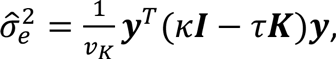, where 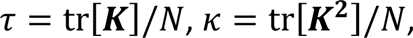, and 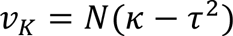. SNP heritability is then estimated as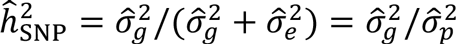.

To estimate the sampling variance of 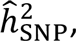 we follow Visscher et al. (2014) and make two assumptions: (1) the off-diagonal elements in the empirical genetic similarity matrix *K* are small, such that *K* ≈ *I* and 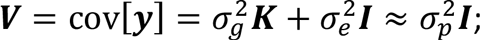 and (2) the phenotypic variance 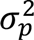 can be estimated with very high precision. We thus have 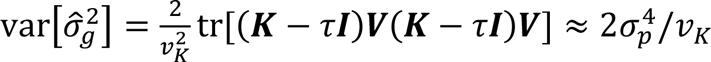, and 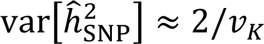. This estimator coincides with existing results in the literature (Visscher et al., 2014). We note that the calculation of the variance of 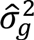 relies on an additional assumption that the trait *y* is Gaussian distributed and thus may be suboptimal for binary traits. However, Visscher and colleagues have empirically shown that this sampling variance approximation is accurate for both continuous and binary traits when the sample size is large (Visscher et al., 2014).

For binary traits, the above calculation gives a heritability estimate on the observed scale, which is dependent on prevalence of the trait in the population. We transformed this heritability estimate to the underlying liability scale under the assumption of a classical liability threshold model (Falconer, 1965; Pearson and Lee, 1901), which makes heritability estimates independent of prevalence and thus comparable across traits. Specifically, heritability estimate on the liability scale can be obtained using a linear transformation of the heritability on the observed scale: 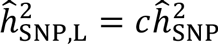, where *c* = *K*(1 — *K*)/φ(*t*)^2^, *K* is the population prevalence, *t* = φ^−1^ (1 – *K*) is the liability threshold, φ is the cumulative distribution function of the standard normal distribution, and *φ* is the density function of the standard normal distribution (Dempster and Lerner, 1950; Lee et al., 2011). Since the UK Biobank is not designed to be ascertained for particular diseases, we assumed that population prevalence is identical to sample prevalence. The sampling variance of the heritability estimate can be transformed accordingly: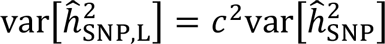.

### Statistical Analysis

In all heritability analyses, we included genotyping array, UK Biobank assessment center, age at recruitment and top 10 principal components (PCs) of the genotype data as covariates. Other covariates such as sex and handedness (e.g., when analyzing the grip strength of the left/right hand) were adjusted where appropriate. See Supplementary Table 1 for the set of covariates we included in the model when estimating the heritability for each trait. To compute PCs of the genotype data, we performed pairwise linkage disequilibrium (LD) based SNP pruning at *R*^2^>0.02 and excluded SNPs in the major histocompatibility complex (MHC) region (chr6:25-35Mb) and chromosome 8 inversion (chr8:7-13Mb). Top PCs were then computed using flashPCA (Abraham and Inouye, 2014) on the pruned data, which employs an efficient randomized algorithm and is thus scalable to large data sets with hundreds of thousands of individuals.

To examine how heritability estimates are modified by sex, we estimated heritability for each non-sex-specific trait in males and females separately. To test if heritability estimates are significantly different by sex, we assumed that the two SNP heritability estimates to be contrasted, *ĥ*_*A*_ and *ĥ*_*B*_, are independent and approximately Gaussian distributed, and computed the z-score of their difference: 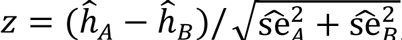, where 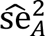 and 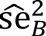 are standard error estimates of *ĥ*_*A*_ and *ĥ*_*B*_, respectively. A *p*-value can then be computed as *p* = 2 · Φ(– |*z*|), where Φ is the cumulative distribution function of the standard normal distribution.

To examine whether SNP heritability estimates vary with age, we used a sliding window approach and estimated heritability for every age range of 10 years (i.e., 40-49 years, 41-50 years, …, 64-73 years) by stratifying samples. We assessed whether heritability estimates exhibited a linear trend with age by fitting a regression model, 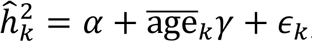, where 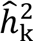 is the heritability estimate in the *k*-th age range, 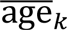 is the mean of the age range, *α* is an intercept, *γ* is the slope and *ε_k_* is the residual of the regression, and testing whether *γ* is significantly different from zero. We weighted heritability estimates by the inverse of their standard errors when fitting the regression model, and thus put more emphasis on estimates with better precision. We only analyzed physical and cognitive measures, and did not consider disease codes and medical history in age stratification analyses because age at recruitment does not reflect disease onset.

Similarly, we used a sliding window approach to estimate the SNP heritability for each trait from the bottom 1/3 quantile to the top 1/3 quantile of the Townsend deprivation index at recruitment, a measure of material deprivation within the population of a given area. For traits that do not reflect the status of participants at the time of recruitment (e.g., medical history and early-life factors), we have implicitly made an assumption that the SES of participants had not changed dramatically throughout their lives.

To account for multiple testing in our stratification analyses, we corrected the *p*-values using the effective number of independent traits we analyzed. Specifically, for each stratification analysis (sex, age and SES), we calculated the Pearson correlation coefficient for each pair of the traits using their overlapping samples. The correlation between traits that had no sample overlap, e.g., male- and female-specific factors, was set to zero. We then conducted a principal component analysis (PCA) to the constructed phenotypic correlation matrix, and estimated the effective numbers of independent traits that explained 99% of the total phenotypic variation in sex, age and SES stratification analyses to be 400, 31 and 440, respectively. Finally, we multiplied uncorrected *p*-values by the corresponding effective number of independent traits to obtain corrected *p*-values.

## Results

We report the heritability for 551 traits that were made available to us through the UK Biobank and had sufficient sample sizes to achieve accurate heritability estimates (standard error of the heritability estimate smaller than 0.1). These traits can be classified into 11 general phenotypic domains: cognitive function (5 traits), early life factors (7 traits), health and medical history (60 traits), hospital in-patient main diagnosis ICD-10 codes (194 traits), life style and environment (88 traits), physical measures (50 traits), psychosocial factors (40 traits), self-reported cancer codes (9 traits), self-reported non-cancer illness codes (79 traits), sex-specific factors (14 traits), and sociodemographics (5 traits). Figure 1 shows the percentage of each domain that makes up the 551 traits we analyzed. Using the top-level categories and chapters of the selfreported disease and ICD-10 coding tree, we can further break down self-reported non-cancer illness codes and ICD-10 codes into different functional domains (Supplementary Figure 2). We note that since we only analyzed disease codes that had prevalence greater than 1% in the sample, distribution of the disease traits across functional domains was highly skewed. For example, we investigated a large number of gastrointestinal and musculoskeletal traits, while diseases that have low prevalence in the sampled population such as psychiatric disorders were not well represented.

**Figure 1:**
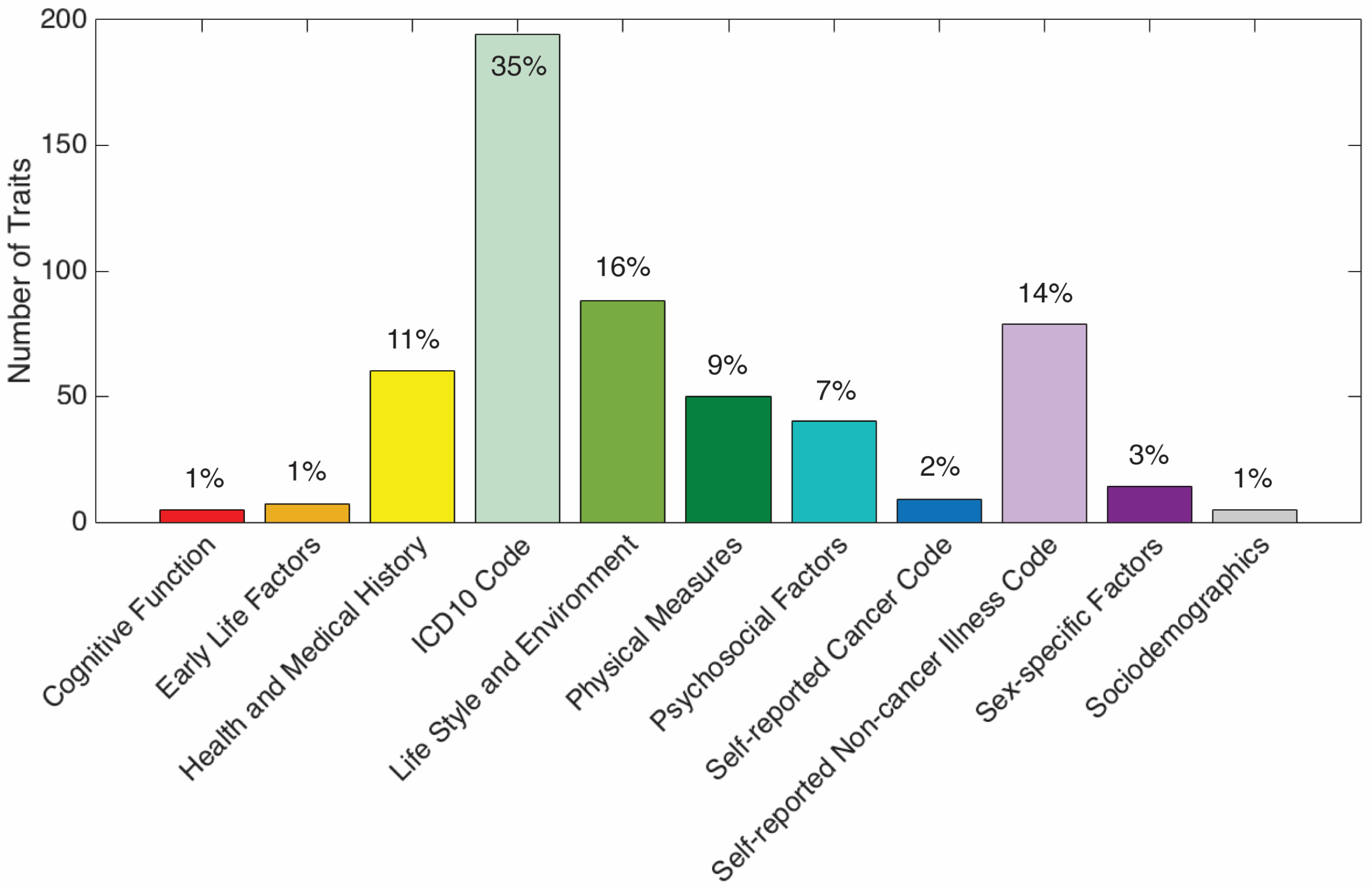
The number of traits in each of the 11 phenotypic domains that make up the 551 traits analyzed in the UK Biobank: cognitive function (5 traits), early life factors (7 traits), health and medical history (60 traits), hospital in-patient main diagnosis ICD-10 codes (194 traits), life style and environment (88 traits), physical measures (50 traits), psychosocial factors (40 traits), self-reported cancer codes (9 traits), selfreported non-cancer illness codes (79 traits), sex-specific factors (14 traits), and sociodemographics (5 traits).

Table 1 lists the top heritable traits in each domain (the most heritable trait and traits with heritability estimates greater than 0.30). Supplementary Tables 1 and 2 show the heritability estimates, standard error estimates, sample sizes, covariates adjusted, prevalence in the sample (for binary traits) and other relevant information for all the traits we analyzed. Common genetic variants appear to have an influence on most traits we investigated, although heritability estimates showed heterogeneity within and across trait domains. Complex traits that exhibited high SNP heritability (larger than 0.40) included human height (0.685+/-0.004), skin color (very fair/fair vs. other, 0.556+/-0.008), ease of skin tanning (very/moderately tanned vs. mildly/occasionally/never tanned, 0.454+/-0.006), comparative height at age 10 (taller than average, 0.439+/-0.007; shorter than average, 0.405+/-0.008), rheumatoid arthritis (0.821+/-0.046), hypothyroidism/myxedema (0.814+/-0.017), malignant neoplasm of prostate (0.426+/-0.093), and diabetes diagnosed by doctor (0.414+/-0.016), among others. On the other end of the spectrum, traits such as duration of walks/moderate activity/vigorous activity, frequency of stair climbing, ever had stillbirth, spontaneous miscarriage or termination, painful gums, stomach disorder, fracture, injuries to the head/knee/leg, and pain in joint had zero or close to zero heritability estimates, indicating that their phenotypic variation is largely determined by environmental factors, or there is widespread heterogeneity or substantial measurement error in these phenotypes. Heritability estimates for several phenotypes,including diseases with known immune-mediated pathogenesis (rheumatoid arthritis, psoriasis, diabetes, hypothyroidism), were markedly reduced when the MHC region was excluded from analysis (Supplementary Table 4), and thus need to be interpreted with caution (see Discussion).

A substantial fraction of the phenotypes we examined were based on self-report illness codes or diagnostic (ICD-10) codes, which may be noisy and have low specificity. However, the SNP heritability estimates for 14 pairs of self-reported illness and ICD-10 codes that represent the same or closely matched diseases were largely consistent and had a Pearson correlation of 0.78 (Table 2), indicating that both phenotypic approaches captured useful variations in these phenotypes.

Heritability analysis stratified by sex identified a number of traits whose heritability showed significant difference in males and females after multiple testing correction (Figure 2). For example, the analyses of diastolic and systolic blood pressure, and self-reported hypertension and high blood pressure provided consistent evidence that the heritability of blood pressure related traits and diseases is significantly higher in females than in males. Adjusting for the current smoking status as a covariate in the model produced virtually identical heritability estimates of these traits and the differences between females and males remained significant.

A majority of physical measures showed decreasing heritability with age (Supplementary Table 3). More specifically, 33 out of 50 physical measures had a significant decreasing trend in heritability estimates after accounting for multiple testing correction (mean slope of the 33 traits -0.0035, i.e., heritability estimates decrease by 3.5 percent per decade). The age-varying SNP heritability estimates and their standard errors for 12 traits that showed both significant slopes and significantly different heritability estimates between the first (40-49 years) and last age range (64-73 years) are shown in Figure 3A.

**Figure 2:**
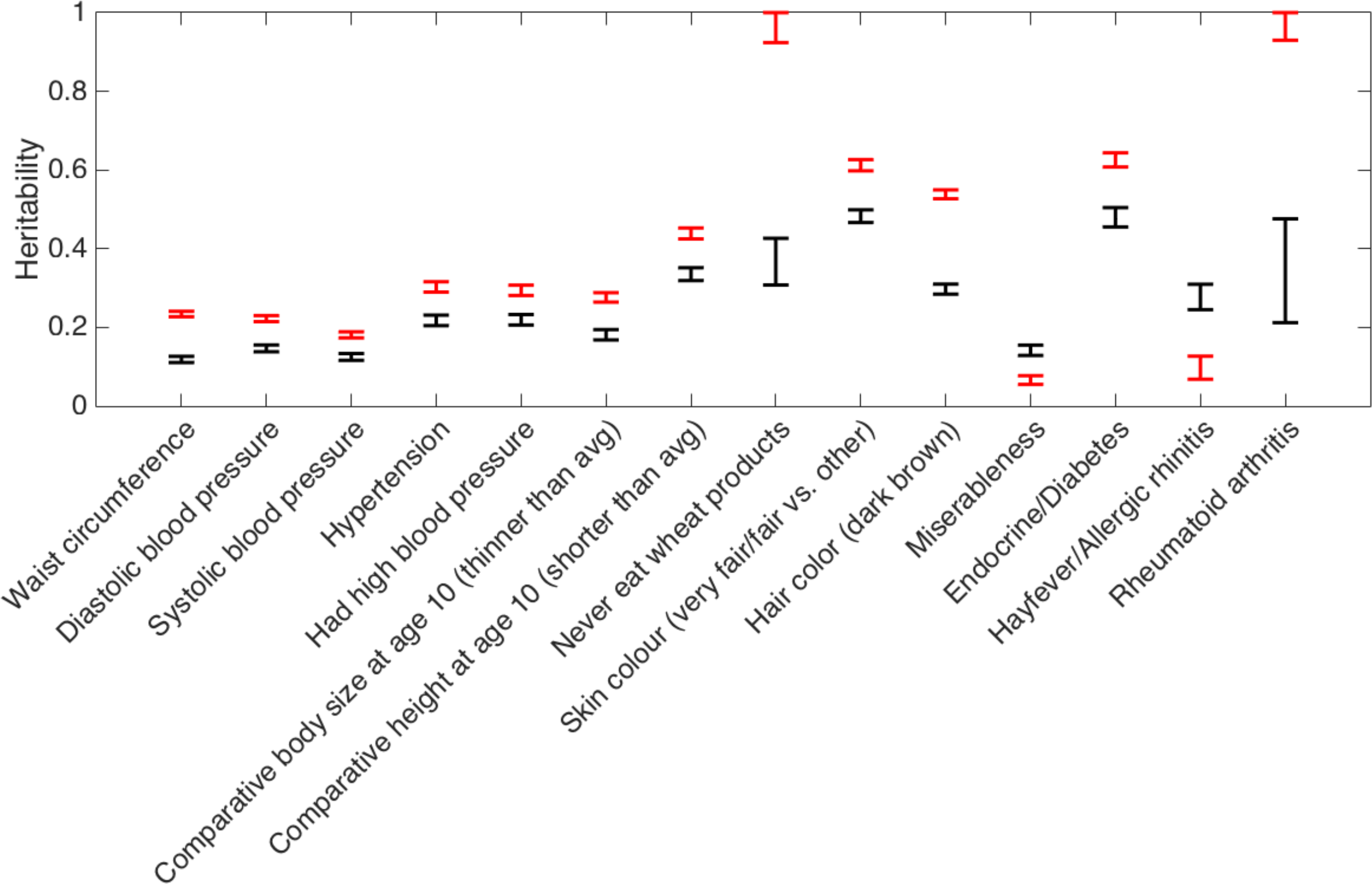
Traits in the UK Biobank that show significantly different SNP heritability estimates in females and males. The heritability estimates of rheumatoid arthritis, endocrine/diabetes and wheat products intake reported here are based on genome-wide SNPs and will be markedly reduced when the major histocompatibility complex (MHC) region (chr6:25-35Mb) is excluded from analysis, and thus need to be interpreted with caution.

When we stratified heritability by the Townsend deprivation index, education (has college or university degree or not) was the only trait on which SES had a significant moderating effect after multiple testing correction. Figure 3B shows that the heritability of education increases with increasing SES.

**Figure 3:**
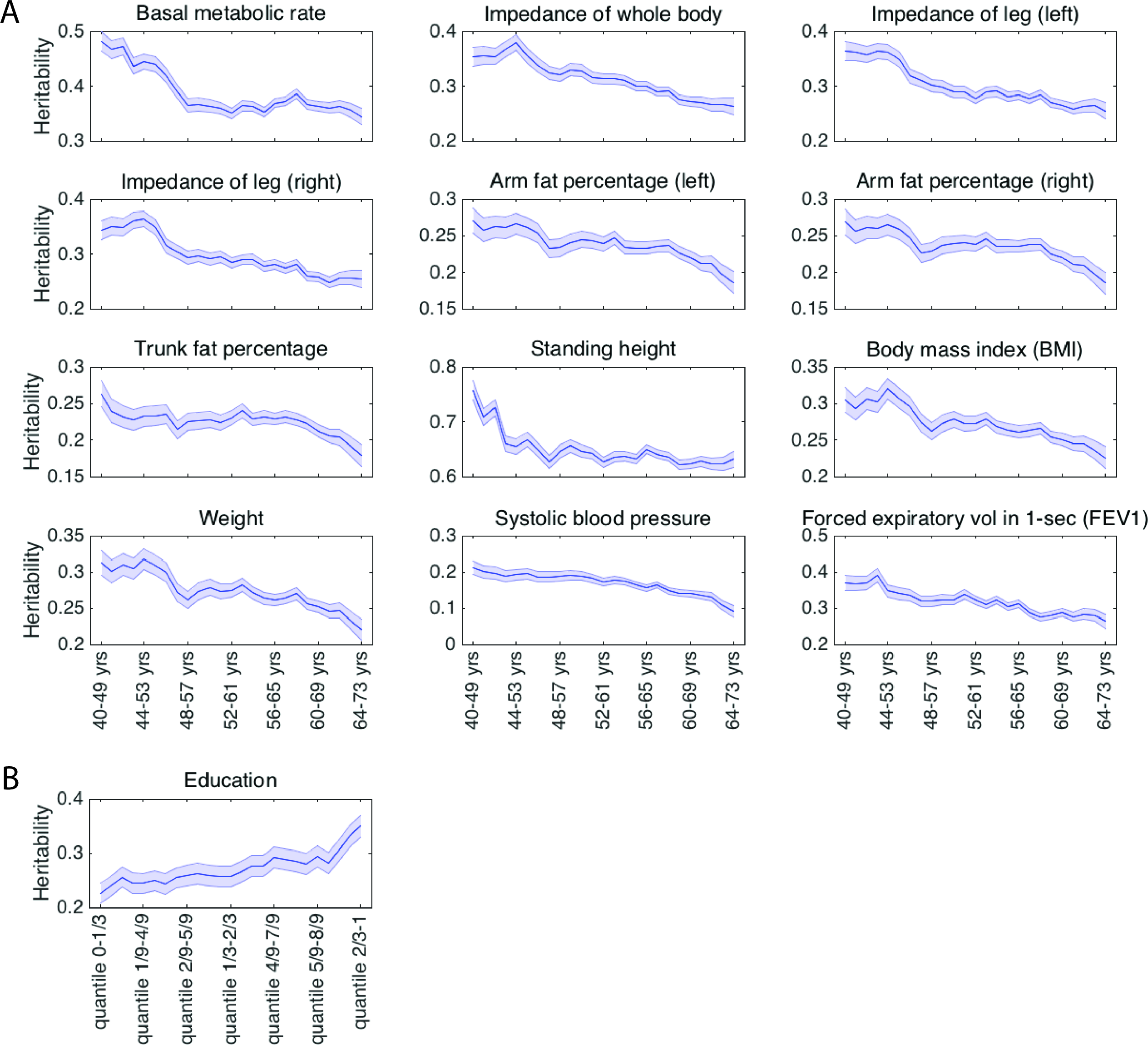
(A) The age-varying heritability estimates and their standard errors (shaded region) for the 12 traits whose heritability significantly decreases with age; (B) The stratified heritability estimates and standard errors (shaded region) of education (has college or university degree or not), on which the socioeconomic status (SES) measured by the Townsend deprivation index has a significant moderating effect.

## Discussion

Estimating the heritability of complex, polygenic traits is an important component of defining the genetic basis of human phenotypes. In addition, heritability estimates provide a theoretical upper bound for the utility of genetic risk prediction models (Chatterjee et al., 2016). We calculated the common-variant heritability of 551 phenotypes derived from the UK Biobank. Three aspects of our work are particularly notable. First, we implemented a computationally efficient method that enabled us to calculate the most extensive population-based survey of SNP heritability to date. Second, we demonstrate that common genetic variation contributes to a broad array of quantitative traits and human diseases in the UK population. Third, we find that the heritability for a number of phenotypes is moderated by major demographic variables, demonstrating the dependence of heritability on population characteristics. We discuss each of these advances below.

Classical methods to estimate SNP heritability rely on the restricted maximum likelihood (ReML) algorithm (Yang et al., 2010, 2011), which can give unbiased heritability estimates in quantitative trait analysis and non-ascertained case-control studies, and is statistically efficient when the trait is Gaussian distributed (Golan et al., 2014). However, ReML is an iterative optimization algorithm whose computational complexity scales cubically with the sample size, and thus can be difficult to apply when analyzing data sets with hundreds of thousands of subjects. In the present study, we used a moment-matching method, which is equivalent to the Haseman-Elston regression (Elston et al., 2000; Haseman and Elston, 1972; Sham and Purcell, 2001) and phenotype-correlation genetic-correlation (PCGC) regression (Golan et al., 2014). The method produces unbiased SNP heritability estimates for both continuous and binary traits, and has much lower computational and memory demand than the ReML algorithm. Once the genetic similarity matrix has been computed, the method takes approximately 20 mins to complete the main heritability analysis and all the stratification analyses for a trait with 100,000 subjects using a single core of the (dual CPU) Intel Xeon 5472 3.0GHz processor and 7 Gb of virtual memory. The moment-matching method is theoretically less statistically efficient than the ReML algorithm (i.e., produces larger standard error on the point estimate) when analyzing quantitative traits, but the power loss is expected to be small (Visscher et al., 2014) and the current sample size is large enough to give accurate heritability estimates for most traits in the UK Biobank.

The moment-matching method is also closely related to the recently developed LD score regression approach, which estimates the SNP heritability of a trait using GWAS summary statistics (Bulik Sullivan et al., 2015b). Specifically, the two methods are mathematically equivalent if (1) the out of sample LD scores estimated from the reference panel and the in sample LD scores estimated from individual-level genotype data are identical; (2) the intercept in the LD score regression model is constrained to 1 (i.e., assuming that there is no confound and population stratification in the data); and (3) a particular weight is used in the LD score regression (more specifically, the reciprocal of the LD score, which is close to the default setting in the LD score regression software) (Bulik Sullivan, 2015). Here, since we have constrained our analysis to white British (Caucasians) and have accounted for potential population stratification by including top PCs of the genotype data as covariates, both methods should produce similar estimates. Compared to LD score regression, our method can avoid conducting a large number of GWAS in stratification analyses and is thus more flexible in this aspect, while LD score regression can be more flexible if we need to meta-analyze heritability estimates from the UK Biobank and other cohorts, partition heritability by functional annotation (Finucane et al., 2015), or estimate the genetic correlation between UK Biobank variables and other complex traits or diseases on which large-scale GWAS results are available (Bulik-Sullivan et al., 2015a; Zheng et al., 2016). To summarize, we have used a method that balances statistical efficiency and computational burden, and is flexible to achieve our main goal in this study.

We found that a large number of traits we examined display significant heritability. For traits whose heritability has been intensively studied, our estimates are generally in line with prior studies. For example, twin and pedigree studies have estimated the heritability of human height and body mass index (BMI) to be approximately 80% and 40-60% (see e.g., Macgregor et al., 2006; Silventoinen et al., 2003, 2008; Visscher et al., 2008), respectively, although recent studies have shown that heritability may be overestimated in family studies due to, for instance, improper modeling of common environment, assortative mating in humans, genetic interactions, and suboptimal statistical methods (Golan et al., 2014; Visscher et al., 2010; Zaitlen et al., 2013, 2014; Zuk et al., 2012). Using genome-wide SNP data from unrelated individuals, it has been shown that common SNPs explain a large proportion of the height and BMI variation in the population, although SNP heritability estimates are lower than twin estimates (Vattikuti et al., 2012; Yang et al., 2010, 2011). Specifically, the first genome-wide complex trait analysis (GCTA, also known as the GREML method) estimated the SNP heritability of human height to be 0.45 using relatively sparse genotyping data (approximately 300,000 SNPs) and showed that the estimate could be higher if imperfect LD between SNPs and causal variants are corrected (Yang et al., 2010). A more recent study leveraging whole-genome sequencing data and imputed genetic variants concluded that narrow-sense heritability is likely to be 60-70% for height and 30-40% for BMI. Here, we estimated the heritability of human height and BMI to be 0.685+/-0.004 and 0.274+/-0.004, respectively, which are comparable to the expected range. Heritability estimates of other complex traits of interest, such as age at menarche in girls (0.239+/-0.007), diastolic (0.184+/-0.004) and systolic (0.156+/-0.004) blood pressures, education (has colleague or university degree or not, 0.294+/-0.007), neuroticism (0.130+/-0.005), smoking (ever smoked or not, 0.174+/-0.006), asthma (0.340+/-0.010) and hypertension (0.263+/-0.007) were also more modest and lower than twin estimates (Poldermann et al., 2015).

Heritability is, by definition, a ratio of variances, reflecting the proportion of phenotypic variance attributable to individual differences in genotypes. Because the genetic architecture and non-genetic influences on a trait may differ depending on the population sampled, heritability itself may vary. Examples of this have been reported in the twin literature. In one well-known study, Turkheimer and colleagues (Turkheimer et al., 2003) reported that the heritability of IQ is moderated by SES in a sample of 320 7-year-old twin pairs of mixed ancestry. In that study, the heritability of IQ was essentially 0 at the lowest end of SES but substantial at the highest end. Subsequent studies of twins at varying ages have produced mixed results (Bates et al., 2013; Hanscombe et al., 2012; Kirkpatrick et al., 2015; Turkheimer et al., 2015). However, in our analysis, using SNP data, we observed no moderating effect of SES (as measured by the Townsend deprivation index) on the heritability of cognitive traits (including fluid intelligence), possibly due to the age range of participants in the UK Biobank (middle and old age) in contrast to many previous studies targeting childhood or early adulthood. On the other hand, the heritability of education did show significant interactions with SES, with increasing heritability at higher SES levels. Prior evidence has suggested that education has substantial genetic correlation with IQ and may be a suitable proxy phenotype for genetic analyses of cognitive performance (Rietveld et al., 2014); thus our results may indirectly support earlier studies of the SES moderation of IQ heritability.

With two exceptions, significant sex differences we observed indicated greater heritability for women compared to men. Our results are consistent with findings from some twin studies but not others. For example, we found that women exhibited significantly greater heritability for measured waist circumference and blood pressure. Twin studies have also reported greater female heritability for waist circumference (Zillikens et al., 2008) but no substantial sex difference in heritability of blood pressure (Hottenga et al., 2005; Polderman et al., 2015). A substantial difference between the heritability of rheumatoid arthritis (RA) in males compared to females was observed, although the MHC region has a large impact on the SNP heritability estimates of autoimmune diseases, and thus this finding needs to be interpreted with caution (see discussion below). While RA is known to be more common in women, a twin analysis found no sex difference in heritability among Finnish and UK twin pairs, though power was limited in that analysis (MacGregor et al., 2000). Intriguingly, greater heritability was observed among men for the personality trait of miserableness, a component of neuroticism, suggesting that environmental factors may be more influential for this trait among women or that measurement error differs by sex.

We examined age effects on heritability for a subset of variables and found that a number of physical measurements indexing body size, adiposity, height, as well as systolic blood pressure and lung function, showed declining heritability with age. Age-related declines in heritability may reflect the cumulative effect of environmental perturbations over the lifespan. Prior twin studies of age effects on the heritability of anthropometric traits in adults have had inconsistent results (Brown et al., 2003; Ortega-Alonso et al., 2009; Schousboe et al., 2004). A recent meta-analysis of 32 twin studies documented a non-monotonic relationship between BMI heritability and age (from childhood to late adulthood), with a peak around age 20 and decline thereafter (Min et al., 2013). An age-related decline in indices of body size may reflect a decreasing contribution of genetically-regulated growth processes over the lifespan. However, our results were restricted to middle and older age groups so that we are unable to assess the entire trajectory of heritability. Some but not all studies have also suggested varying or declining heritability with age for blood pressure and lung function, as we found here (Brown et al., 2003; Coultas et al., 1991; Hottenga et al., 2006; McClearn et al., 1994; Menni et al., 2013; Vinck et al., 2001; Wang et al., 2015).

Our results should be interpreted in light of several limitations. First, the phenotypes were limited to those for which we had sufficient data to estimate heritability with adequate precision. Therefore, diseases with low prevalence in the sampled population were not well represented in our analysis. We also assumed in our analysis that the population prevalence of a binary trait is identical to the observed sample prevalence, but diseases such as schizophrenia and stroke are naturally under-ascertained and thus their sample prevalence is often lower than population prevalence. In addition, we note that since we used medical history to define cases and controls, the prevalence of many diseases we investigated reflected lifetime prevalence, which may be different from cross-sectional prevalence used in other studies. Second, a substantial fraction of the phenotypes we examined were based on self-report or diagnostic (ICD-10) codes, which may or may not validly capture the phenotypes they represent, although a head-to-head comparison of the heritability estimates between self-reported illness and ICD-10 codes showed largely consistent results. Prior research evaluating phenotypes derived from electronic health records indicate that greater phenotypic validity can be achieved when diagnostic codes are supplemented with text mining methods (Castro et al., 2015; Ford et al., 2016; Liao et al., 2015; Perlis et al., 2012). The specificity of the disease codes may also be improved by leveraging the medication records in the UK Biobank. Third, we binarized categorical (multinomial or ordinal) variables to facilitate analysis, but this might not optimally represent variation in these variables with respect to heritability. Fourth, as heritability can be population specific, our estimates may not generalize to other settings or ancestry groups. Finally, heritability estimation always relies on a number of assumptions on the genetic architecture. For example, the method we used here, as well as the established GCTA and LD score regression approaches, implicitly assumes that the causal SNPs are randomly spread over the genome, which is independent of the MAF spectrum and the LD structure, and the effect sizes of causal SNPs are Gaussian distributed and have a specific relationship to their MAFs. Although it has been shown that SNP heritability estimates are reasonably robust to many of these modeling assumptions (Speed et al., 2012), the estimates can be biased if, for instance, causal SNPs are rarer or more common than uniformly distributed on the MAF spectrum, or are enriched in high or low LD regions across the genome. For example, the heritability estimates for some autoimmune diseases such as psoriasis and rheumatoid arthritis dropped dramatically when the MHC region (chr6:25-35Mb) was removed when constructing the genetic similarity matrix, indicating, as expected, that causal variants for these diseases are disproportionally enriched in the MHC region. Supplementary Table 4 lists all the traits whose heritability estimates decreased by 0.2 or more when the MHC region was taken out, and thus need to be interpreted with caution. Methods to correct for MAF properties and region-specific LD heterogeneity of causal variants have been proposed (Lee et al., 2013; Speed et al., 2012; Yang et al., 2015). For example, we can stratify MAF and LD structure into different bins, compute a genetic similarity matrix within each bin, and fit a mixed effects model with multiple variance components (Lee et al., 2013; Yang et al., 2015). This approach can give heritability estimates that are robust to properties of the underlying genetic architecture, but has the downside of increased computational burden and reduced statistical power. A different direction to explore is to estimate SNP heritability using imputed data (in contrast to the genotype data here), which might capture more genetic variation from rare variants, or common variants that are not well tagged by the genotyped SNPs, and thus lead to increased heritability estimates. Here, as the first study to screen all UK Biobank variables and provide overview on the distribution of SNP heritability across different trait domains, and to examine the effect of potential modifying variables on heritability estimates, we used a straightforward and classical modeling approach that is most widely used. To obtain more insights into the genetic architecture and find the most appropriate model for each individual trait, more systematic investigation is needed.

In sum, using an efficient computational approach, we provide estimates of the SNP heritability for 551 complex traits across the phenome captured in the population-based UK Biobank. We further identify phenotypes for which the contribution of genetic variation is modified by demographic factors. These results underscore the importance of considering population characteristics in comparing heritability, highlight phenotypes and subgroups that may warrant priority for genetic association studies, and may inform efforts to apply genetic risk prediction models for a broad range of human phenotypes.

## Acknowledgements

This research was carried out in part at the Athinoula A. Martinos Center for Biomedical Imaging at the Massachusetts General Hospital (MGH), using resources provided by the Center for Functional Neuroimaging Technologies,P41EB015896, a P41 Biotechnology Resource Grant supported by the National Institute of Biomedical Imaging and Bioengineering (NIBIB), National Institutes of Health (NIH). This work also involved the use of instrumentation supported by the NIH Shared Instrumentation Grant Program; specifically, grant numbers S10RR023043 and S10RR023401. This research was also funded in part by NIH grants R01NS083534, R01NS070963, 1K25EB013649-01 and 1R21AG050122-01A1 (to MRS); K24MH094614 (to JWS); and an MGH ECOR Tosteson Postdoctoral Fellowship Award (to TG). JWS is a Tepper Family MGH Research Scholar and was also supported in part by a gift from the Demarest Lloyd Foundation. This research has been conducted using the UK Biobank resource.

## Tables and Figures

**Tables 1:**The heritability estimates, standard error estimates, sample sizes, covariates adjusted, prevalence in the sample (for binary traits) and other relevant information for the top heritable traits in each phenotypic domain.

**Tables 2:**A head-to-head comparison of SNP heritability estimates for the self-reported illness codes and ICD-10 codes that represent the same or closely matched diseases.

## References

Abraham, G., & Inouye, M. (2014). Fast principal component analysis of large-scale genome-wide data. PLoS One, 9(4), e93766.

Bates, T. C., Lewis, G. J., & Weiss, A. (2013). Childhood socioeconomic status amplifies genetic effects on adult intelligence. Psychological Science, 24(10), 2111–2116.

Brown, W. M., Beck, S. R., Lange, E. M., Davis, C. C., Kay, C. M., Langefeld, C. D., & Rich, S. S.(2003). Age-stratified heritability estimation in the Framingham Heart Study families. BMC Genetics, 4(1), 1.

Bulik-Sullivan, B. (2015). Relationship between LD score and Haseman-Elston regression. bioRxiv, 018283.

Bulik-Sullivan, B., Finucane, H. K., Anttila, V., Gusev, A., Day, F. R., Loh, P. R.,... & Daly, M. J. (2015a). An atlas of genetic correlations across human diseases and traits. Nature Genetics, 47(11), 1236–1241.

Bulik-Sullivan, B. K., Loh, P. R., Finucane, H. K., Ripke, S., Yang, J., Patterson, N.,... & Schizophrenia Working Group of the Psychiatric Genomics Consortium. (2015b). LD Score regression distinguishes confounding from polygenicity in genome-wide association studies. Nature Genetics, 47(3), 291–295.

Castro, V. M., Minnier, J., Murphy, S. N., Kohane, I., Churchill, S. E., Gainer, V.,... & Weill, S. R. (2015). Validation of electronic health record phenotyping of bipolar disorder cases and controls. American Journal of Psychiatry, 172(4), 363–372.

Chang, C. C., Chow, C. C., Tellier, L. C., Vattikuti, S., Purcell, S. M., & Lee, J. J. (2015). Second-generation PLINK: rising to the challenge of larger and richer datasets. Gigascience, 4(1), 1.

Chatterjee, N., Shi, J., & García-Closas, M. (2016). Developing and evaluating polygenic risk prediction models for stratified disease prevention. Nature Reviews Genetics, 17(7), 392–406.

Coultas, D. B., Hanis, C. L., Howard, C. A., Skipper, B. J., & Samet, J. M. (1991). Heritability of ventilatory function in smoking and nonsmoking New Mexico Hispanics. American Review of Respiratory Disease, 144(4), 770–775.

Dempster, E. R., & Lerner, I. M. (1950). Heritability of threshold characters. Genetics, 35(2), 212.

Elston, R. C., Buxbaum, S., Jacobs, K. B., & Olson, J. M. (2000). Haseman and Elston revisited. Genetic Epidemiology, 19(1), 1–17.

Finucane, H. K., Bulik-Sullivan, B., Gusev, A., Trynka, G., Reshef, Y., Loh, P. R., ... & Ripke, S. (2015). Partitioning heritability by functional annotation using genome-wide association summary statistics. Nature Genetics, 47(11), 1228–1235.

Ford, E., Carroll, J.A., Smith, H. E., Scott, D., & Cassell, J. A. (2016). Extracting information from the text of electronic medical records to improve case detection: a systematic review. Journal of the American Medical Informatics Association, ocv180.

Falconer, D. S. (1965). The inheritance of liability to certain diseases, estimated from the incidence among relatives. Annals of Human Genetics, 29(1), 51–76.

Ge, T., Nichols, T. E., Lee, P. H., Holmes, A. J., Roffman, J. L., Buckner, R. L., ... & Smoller, J. W. (2015). Massively expedited genome-wide heritability analysis (MEGHA). Proceedings of the National Academy of Sciences, 112(8), 2479–2484.

Golan, D., Lander, E. S., & Rosset, S. (2014). Measuring missing heritability: inferring the contribution of common variants. Proceedings of the National Academy of Sciences, 111(49), E5272–E5281.

Hanscombe, K. B., Trzaskowski, M., Haworth, C. M., Davis, O. S., Dale, P. S., & Plomin, R. (2012). Socioeconomic status (SES) and children's intelligence (IQ): In a UK-representative sample SES moderates the environmental, not genetic, effect on IQ. PLoS One, 7(2), e30320.

Haseman, J. K., & Elston, R. C. (1972). The investigation of linkage between a quantitative trait and a marker locus. Behavior Genetics, 2(1), 3–19.

Hottenga, J. J., Boomsma, D. I., Kupper, N., Posthuma, D., Snieder, H., Willemsen, G., & de Geus, E. J. (2005). Heritability and stability of resting blood pressure. Twin Research and Human Genetics, 8(05), 499–508.

Hottenga, J. J., Whitfield, J. B., De Geus, E. J., Boomsma, D. I., & Martin, N. G. (2006). Heritability and stability of resting blood pressure in Australian twins. Twin Research and Human Genetics, 9(02), 205–209.

Kirkpatrick, R. M., McGue, M., & Iacono, W. G. (2015). Replication of a gene-environment interaction via multimodel inference: additive-genetic variance in Adolescents' General Cognitive Ability Increases with Family-of-Origin Socioeconomic Status. Behavior Genetics, 45(2), 200–214.

Lee, S. H., Wray, N. R., Goddard, M. E., & Visscher, P. M. (2011). Estimating missing heritability for disease from genome-wide association studies. The American Journal of Human Genetics, 88(3), 294–305.

Lee, S. H., Yang, J., Chen, G. B., Ripke, S., Stahl, E. A., Hultman, C. M., ... & Wray, N. R. (2013). Estimation of SNP heritability from dense genotype data. The American Journal of Human Genetics, 93(6), 1151–1155.

Liao, K. P., Cai, T., Savova, G. K., Murphy, S. N., Karlson, E. W., Ananthakrishnan, A. N.,... & Churchill, S. (2015). Development of phenotype algorithms using electronic medical records and incorporating natural language processing. BMJ, 350, h1885.

Pearson, K., & Lee, A. (1901). Mathematical contributions to the theory of evolution VII — On the application of certain formulae in the theory of correlation to the inheritance of characters not capable of quantitative measurement. Proceedings of the Royal Society of London, 66(424–433), 324–327.

MacGregor, S., Cornes, B. K., Martin, N. G., & Visscher, P. M. (2006). Bias, precision and heritability of selfreported and clinically measured height in Australian twins. Human Genetics, 120(4), 571–580.

MacGregor, A. J., Snieder, H., Rigby, A. S., Koskenvuo, M., Kaprio, J., Aho, K., & Silman, A. J. (2000). Characterizing the quantitative genetic contribution to rheumatoid arthritis using data from twins. Arthritis & Rheumatism, 43(1), 30.

McClearn, G. E., Svartengren, M., Pedersen, N. L., Heller, D. A., & Plomin, R. (1994). Genetic and environmental influences on pulmonary function in aging Swedish twins. Journal of Gerontology, 49(6), M264–M268.

Menni, C., Mangino, M., Zhang, F., Clement, G., Snieder, H., Padmanabhan, S., & Spector, T. D. (2013). Heritability analyses show visit-to-visit blood pressure variability reflects different pathological phenotypes in younger and older adults: evidence from UK twins. Journal of Hypertension, 31(12), 2356–2361.

Min, J., Chiu, D. T., & Wang, Y. (2013). Variation in the heritability of body mass index based on diverse twin studies: a systematic review. Obesity Reviews, 14(11), 871–882.

Ortega-Alonso, A., Sipilä, S., Kujala, U. M., Kaprio, J., & Rantanen, T. (2009). Genetic influences on change in BMI from middle to old age: a 29-year follow-up study of twin sisters. Behavior Genetics, 39(2), 154–164.

Perlis, R. H., Iosifescu, D. V., Castro, V. M., Murphy, S. N., Gainer, V. S., Minnier, J., ... & Fava, M. (2012). Using electronic medical records to enable large-scale studies in psychiatry: treatment resistant depression as a model. Psychological Medicine, 42(1), 41–50.

Polderman, T. J., Benyamin, B., De Leeuw, C. A., Sullivan, P.F., Van Bochoven, A., Visscher, P.M., & Posthuma, D. (2015). Meta-analysis of the heritability of human traits based on fifty years of twin studies. Nature Genetics, 47(7), 702–709.

Rietveld, C.A., Esko, T., Davies, G., Pers, T.H., Turley, P., Benyamin, B., ... & De Leeuw, C. (2014). Common genetic variants associated with cognitive performance identified using the proxy-phenotype method. Proceedings of the National Academy of Sciences, 111(38), 13790–13794.

Schousboe, K., Visscher, P.M., Erbas, B., Kyvik, K.O., Hopper, J.L., Henriksen, J.E., ... & Sørensen, T. I. A. (2004). Twin study of genetic and environmental influences on adult body size, shape, and composition. International Journal of Obesity, 28(1), 39–48.

Sham, P.C., & Purcell, S. (2001). Equivalence between Haseman-Elston and variance-components linkage analyses for sib pairs. The American Journal of Human Genetics, 68(6), 1527–1532.

Silventoinen, K., Magnusson, P.K., Tynelius, P., Kaprio, J., & Rasmussen, F. (2008). Heritability of body size and muscle strength in young adulthood: a study of one million Swedish men. Genetic Epidemiology, 32(4), 341–349.

Silventoinen, K., Sammalisto, S., Perola, M., Boomsma, D.I., Cornes, B.K., Davis, C., ... & Luciano, M. (2003). Heritability of adult body height: a comparative study of twin cohorts in eight countries. Twin Research, 6(05), 399–408.

Speed, D., Hemani, G., Johnson, M.R., & Balding, D. J. (2012). Improved heritability estimation from genome-wide SNPs. The American Journal of Human Genetics, 91(6), 1011–1021.

Sudlow, C., Gallacher, J., Allen, N., Beral, V., Burton, P., Danesh, J., ... & Liu, B. (2015). UK biobank: an open access resource for identifying the causes of a wide range of complex diseases of middle and old age. PLoS Medicine, 12(3), e1001779.

Turkheimer, E., Beam, C.E., & Davis, D. W. (2015). The Scarr-Rowe interaction in complete seven-year WISC data from the Louisville twin study: Preliminary report. Behavior Genetics, 45(6), 635–639.

Turkheimer, E., Haley, A., Waldron, M., D'Onofrio, B., & Gottesman, I. I. (2003). Socioeconomic status modifies heritability of IQ in young children. Psychological Science, 14(6), 623–628.

Vattikuti, S., Guo, J., & Chow, C. C. (2012). Heritability and genetic correlations explained by common SNPs for metabolic syndrome traits. PLoS Genetics, 8(3), e1002637.

Vinck, W.J., Fagard, R.H., Loos, R., & Vlietinck, R. (2001). The impact of genetic and environmental influences on blood pressure variance across age-groups. Journal of Hypertension, 19(6), 1007–1013.

Visscher, P.M., Hemani, G., Vinkhuyzen, A.A., Chen, G.B., Lee, S.H., Wray, N.R., ... & Yang, J. (2014). Statistical power to detect genetic (co) variance of complex traits using SNP data in unrelated samples. PLoS Genetics, 10(4), e1004269.

Visscher, P.M., Hill, W.G., & Wray, N. R. (2008). Heritability in the genomics era—concepts and misconceptions. Nature Reviews Genetics, 9(4), 255–266.

Visscher, P.M., McEvoy, B., & Yang, J. (2010). From Galton to GWAS: quantitative genetics of human height. Genetics Research, 92, 371–379.

Wang, B., Liao, C., Zhou, B., Cao, W., Lv, J., Yu, C., ... & Li, L. (2015). Genetic contribution to the variance of blood pressure and heart rate: A systematic review and meta-regression of twin studies. Twin Research and Human Genetics, 18(02), 158–170.

Yang, J., Bakshi, A., Zhu, Z., Hemani, G., Vinkhuyzen, A.A., Lee, S.H., ... & Snieder, H. (2015). Genetic variance estimation with imputed variants finds negligible missing heritability for human height and body mass index. Nature Genetics, 47(10), 1114–1120.

Yang, J., Benyamin, B., McEvoy, B.P., Gordon, S., Henders, A.K., Nyholt, D.R., ... & Goddard, M. E. (2010). Common SNPs explain a large proportion of the heritability for human height. Nature Genetics, 42(7), 565–569.

Yang, J., Lee, S.H., Goddard, M.E., & Visscher, P. M. (2011). GCTA: a tool for genome-wide complex trait analysis. The American Journal of Human Genetics, 88(1), 76–82.

Zaitlen, N., Kraft, P., Patterson, N., Pasaniuc, B., Bhatia, G., Pollack, S., & Price, A. L. (2013). Using extended genealogy to estimate components of heritability for 23 quantitative and dichotomous traits. PLoS Genetics, 9(5), e1003520.

Zaitlen, N., Pasaniuc, B., Sankararaman, S., Bhatia, G., Zhang, J., Gusev, A., ... & Assimes, T. L. (2014). Leveraging population admixture to characterize the heritability of complex traits. Nature Genetics, 46(12), 1356–1362.

Zheng, J., Erzurumluoglu, M., Elsworth, B., Howe, L., Haycock, P., Hemani, G., ... & Finucane, H. (2016). LD Hub: a centralized database and web interface to perform LD score regression that maximizes the potential of summary level GWAS data for SNP heritability and genetic correlation analysis. bioRxiv, 051094.

Zillikens, M.C., Yazdanpanah, M., Pardo, L.M., Rivadeneira, F., Aulchenko, Y.S., Oostra, B.A., ... & van Duijn, C. M. (2008). Sex-specific genetic effects influence variation in body composition. Diabetologia, 51(12), 2233–2241.

Zuk, O., Hechter, E., Sunyaev, S.R., & Lander, E. S. (2012). The mystery of missing heritability: Genetic interactions create phantom heritability. Proceedings of the National Academy of Sciences, 109(4),1193–1198.

